# What scRNA sequencing taught us about MGMT expression in glioblastoma

**DOI:** 10.1101/2024.07.23.604798

**Authors:** Iyad Alnahhas, Mehak Khan, Wenyin Shi

**Affiliations:** Division of Neuro-Oncology, Department of Neurology, Thomas Jefferson University; Sidney Kimmel Medical College, Thomas Jefferson University; Department of Radiation Oncology, Thomas Jefferson University

## Abstract

**Introduction:** The promoter methylation status of O-6-methylguanine-DNA methyltransferase (MGMTp) is an established predictive and prognostic marker in GBM. Previous studies showed that the expression of MGMT based on immunohistochemistry was variable and lacked association with survival. This in part is because non-tumor cells including endothelial cells and macrophages express MGMT. Advanced technologies such as single-cell RNA (scRNA) sequencing have helped to elucidate the cellular composition of cancer and its microenvironment. scRNA sequencing allows to assess gene expression level in tumor cells specifically.

**Methods:** We used publicly available data from two recent GBM scRNA studies that included MGMTp methylation status data for patients to explore and uncover details about MGMT expression at the single-cell level: CPTAC (13 primary samples) and Neftel (20 primary samples).

**Results:** In the CPTAC study, MGMT expression ranged from 0.19%-1.43% in the MGMTp methylated group (median 0.82%), and from 2.17%-28.36% in the MGMTp unmethylated group (median 5.7%). It therefore appears that 2% is a reasonable expression cutoff to predict the MGMTp methylation status based on scRNA data. In the Neftel study, MGMT expression ranged from 0-1.26% in the MGMTp methylated group (median 0.59%), and from 0.3-27.67% in the MGMTp unmethylated group (median 12.44%). Three unmethylated samples (out of 16) did not follow the 2% rule. It remains unclear if this is due to technical inaccuracies as the Neftel paper did not specify the method used to detect MGMTp methylation or even mere typos. Alternatively, could it be that truly MGMTp unmethylated samples can have low MGMT expression? Could this explain why some unmethylated MGMTp GBM patients surpass the expected survival? Interestingly, gene set enrichment analysis shows that MGMT expressing cells are enriched with mesenchymal genes, whereas MGMT negative cells are enriched with proneural genes.

**Conclusion:** Fewer than 2% of GBM cells express MGMT when MGMTp is methylated.

## Introduction

Astrocytomas in adults are classified into isocitrate dehydrogenase (IDH) mutant and IDH-wild-type (IDHwt) subtypes. IDH mutant astrocytomas more commonly occur in younger patients and carry a relatively better prognosis. IDHwt astrocytomas, on the other hand, more commonly occur in older patients and carry worse prognosis. Glioblastoma (GBM) represents IDHwt astrocytoma grade IV (1). The standard of care for GBM includes maximal safe resection followed by concurrent radiotherapy with an oral alkylating agent (temozolomide) and adjuvant temozolomide (2). The promoter methylation status of O-6-methylguanine-DNA methyltransferase (MGMTp) is the most established molecular predictive marker for response to temozolomide and accordingly impacts overall survival in GBM (3). Previous literature suggests that the median overall survival (OS) for patients with unmethylated MGMTp GBM is 14.11 months with a median progression-free survival (PFS) of 4.99 months. In contrast, the median OS for patients with methylated MGMTp GBM is 24.59 months with a PFS of 9.51 months (4).

Despite the significance of MGMTp methylation status on survival in GBM, previous studies showed that the expression of MGMT based on immunohistochemistry (IHC) in GBM samples was variable and lacked association with survival (5, 6). This is in part because non-tumor cells including endothelial cells and macrophages can express MGMT limiting accurate interpretations of the IHC stains. Furthermore, it has been observed that the MGMTp methylation status can change between paired primary and recurrent samples in 19-24% of cases, more commonly from methylated to unmethylated status but also the other way around. It appears that the MGMTp methylation status in the recurrence setting is of less significance on survival (7, 8).

Advanced technologies such as single-cell RNA (scRNA) sequencing and spatial transcriptomics have helped to elucidate the cell-type composition of cancer and its microenvironment. Recent scRNA/single-nucleus RNA sequencing (snRNA-seq) studies have demonstrated that GBM cells exhibit a high degree of heterogeneity and plasticity and seamless transitions between cellular states (9, 10).

In this paper, we use publicly available data from three recent scRNA/snRNA IDHwt GBM studies (9-11) to explore and uncover details about MGMT expression in GBM in light of MGMTp methylation status at the single-cell level.

## Methods

We first used snRNA data from 18 treatment-naive GBM patients prospectively collected by the Clinical Proteomic Tumor Analysis Consortium (11). The cohort is well annotated and includes information about the MGMT promoter methylation status for samples as determined by the MGMT-STP27 model from DNA methylation data. Data was downloaded from the GDC Data Portal. Details about the files used in this analysis can be found in the supplementary document. Cell-type annotation (tumor cells versus microenvironment) were applied per the original paper’s annotation.

We then aimed to validate the above findings by evaluating a different dataset by Neftel et al (9). The study performed scRNA sequencing on 20 adult IDHwt GBM samples. The supplementary table was downloaded from the original paper and included the clinical characteristics for the cohort including the MGMT promoter methylation status. However, the MGMTp determination method was not specified in the paper. The pre-processed matrix file was downloaded from the 3CA database (12). The 3CA database houses 77 scRNA datasets where the quality control, filtering and cell-type annotation were all consistently applied to all the datasets and made available to download.

We finally applied the findings to the study by Wang, et al (10). The study profiled 86 primary-recurrent patient-matched paired GBM specimens with snRNA sequencing. 76 of these samples were IDHwt GBM. The study did not include MGMTp methylation data for the samples. However, we were interested in the change of MGMT expression between the primary and recurrent samples. The data was downloaded from GEO using the accession number GSE174554.

We used R 4.3.1 to analyze the scRNA/snRNA datasets. All code used is available on GitHub (https://github.com/iyadalnahhas/scRNA_MGMT/blob/main/scRNA_MGMT.Rmd)

Seurat objects were created for the above studies per the Seurat V5 workflow (13). For the CPTAC and Wang et al studies, quality control was completed as follows: cells were selected for further analysis after excluding potential empty droplets (less than 200 genes per cell) and two few unique molecular identifiers (UMIs <1000) and doublets or multiplets (cells with more than 10000 genes per cell). Low quality or dying cells were excluded by selecting cells with less than 10% mitochondrial genes. The data was then normalized and highly variable features were selected. The data was then scaled and dimensionality reductions were applied.

MGMT expression was determined using the FetchData command from Seurat. Cell-cycle scores were calculated, and cell-cycle classification predictions (G2M, S or G1 phase) were applied per the Seurat workflow. We then used Seurat’s FindMarkers function to find the differentially expressed genes between the cellular groups of interest. As a default, Seurat uses the non-parameteric Wilcoxon rank sum test to perform this analysis. Gene set enrichment analysis was performed using clusterProfiler (14). The C2 curated gene collection set was used from the Molecular Signatures Database (MSigDB) using the msigdbr package in R.

## Results

### Percentages of MGMT expression per MGMTp status

#### # CPTAC study

MGMTp methylation data for the CPTAC snRNA cohort was available for 13 patients (7 patients with unmethylated MGMT and 6 patients with methylated MGMT) whose samples were included in this analysis. The number of tumor cells per sample after quality control ranged from 1121-10996 (median 5447 cells). MGMT expression data was extracted by Seurat and cells were classified into MGMT expressing (MGMT+) and MGMT not-expressing (MGMT-).

Table 1 shows MGMT expression for each of the 13 samples in this cohort. MGMT expression ranged from 0.19%-1.43% in the MGMTp methylated group (median 0.82%), and from 2.17%-28.36% in the MGMTp unmethylated group (median 5.7%). It therefore appears that 2% cellular expression represents a reasonable cutoff to predict the MGMTp methylation status based on scRNA expression data.

**Table 1:** Percentage of MGMT expression per sample in the CPTAC cohort.

| Case ID | MGMTp status | MGMT expression % |
| --- | --- | --- |
| C3N-03184 | Methylated | 0.19% |
| C3N-02181 | Methylated | 0.71% |
| C3L-03968 | Methylated | 0.77% |
| C3L-02705 | Methylated | 0.88% |
| C3N-02769 | Methylated | 1.35% |
| C3L-03405 | Methylated | 1.43% |
| C3N-01798 | Unmethylated | 2.17% |
| C3N-02784 | Unmethylated | 4.01% |
| C3N-02783 | Unmethylated | 4.89% |
| C3N-03186 | Unmethylated | 5.7% |
| C3N-03188 | Unmethylated | 8.37% |
| C3N-02190 | Unmethylated | 9.43% |
| C3N-01816 | Unmethylated | 28.36% |

#### # Neftel et al study

MGMT promoter methylation data for the Neftel scRNA cohort was available for 20 patients (16 patients with unmethylated MGMT and 4 patients with methylated MGMT). The number of tumor cells per sample after quality control ranged from 121-435 (median 221 cells). MGMT expression data was extracted by Seurat and cells were classified into MGMT expressing (MGMT+) and MGMT not-expressing (MGMT-).

Table 2 shows MGMT expression for each of the 20 samples in this cohort. MGMT expression ranged from 0-1.26% in the MGMTp methylated group (median 0.59%), and from 0.3-27.67% in the MGMTp unmethylated group (median 12.44%). Three unmethylated samples (out of 16) did not follow the rule of MGMT+ cells >2% (0.3%, 0.68%, 0.73%). Therefore, the <2% cutoff rule has 100% sensitivity and 81.25% specificity to predict unmethylated MGMTp status.

**Table 2:** Percentage of MGMT expression per sample in the Neftel et al cohort.

| Case ID | MGMTp status | MGMT expression % |
| --- | --- | --- |
| MGH152 | Methylated | 0% |
| MGH66 | Methylated | 0.23% |
| MGH100 | Methylated | 0.96% |
| MGH128 | Methylated | 1.26% |
| MGH125 | Unmethylated | 0.3% |
| MGH121 | Unmethylated | 0.68% |
| MGH143 | Unmethylated | 0.96% |
| MGH101 | Unmethylated | 3.92% |
| MGH115 | Unmethylated | 4.38% |
| MGH124 | Unmethylated | 7.34% |
| MGH105 | Unmethylated | 9.6% |
| MGH106 | Unmethylated | 11.76% |
| MGH151 | Unmethylated | 14.05% |
| MGH129 | Unmethylated | 15.79% |
| MGH110 | Unmethylated | 18.27% |
| MGH104 | Unmethylated | 19.44% |
| MGH122 | Unmethylated | 21% |
| MGH113 | Unmethylated | 21.3% |
| MGH136 | Unmethylated | 23.14% |
| MGH102 | Unmethylated | 27.67% |

#### # Wang et al study

Of the 76 IDHwt samples, 48 were paired (primary/recurrent) and passed the above specified quality control. We excluded samples having fewer than 10 malignant cells after quality control. This left 42 samples (21 pairs). The number of tumor cells per sample ranged from 12-5868 (median 666.5 cells). MGMT expression data was extracted by Seurat and cells were classified into MGMT expressing (MGMT+) and MGMT not-expressing (MGMT-).

Of the 21 paired samples, MGMT expression decreased at recurrence in 11 pairs and increased in 10 pairs. By using the 2% cutoff, MGMTp methylation status changed from unmethylated to methylated in 4/21 pairs (19%) and from methylated to unmethylated in 4/21 pairs (19%). Figure 1 shows a ladder plot demonstrating the change in MGMT expression percentage between primary and recurrent samples.

**Figure 1:**
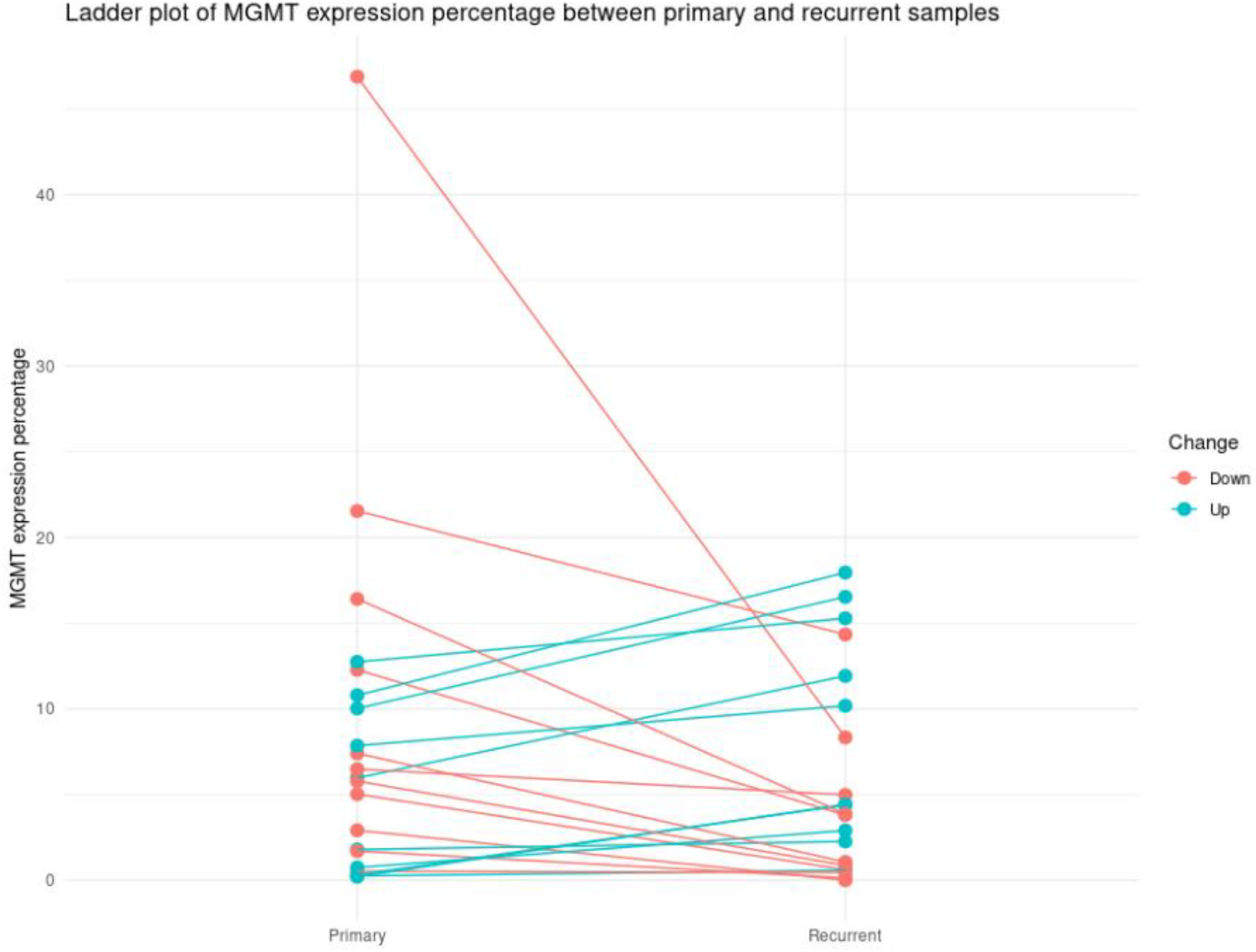
A ladder plot showing the change in MGMT expression percentage between primary and recurrent samples in the Wang et al cohort.

**Figure 2:**
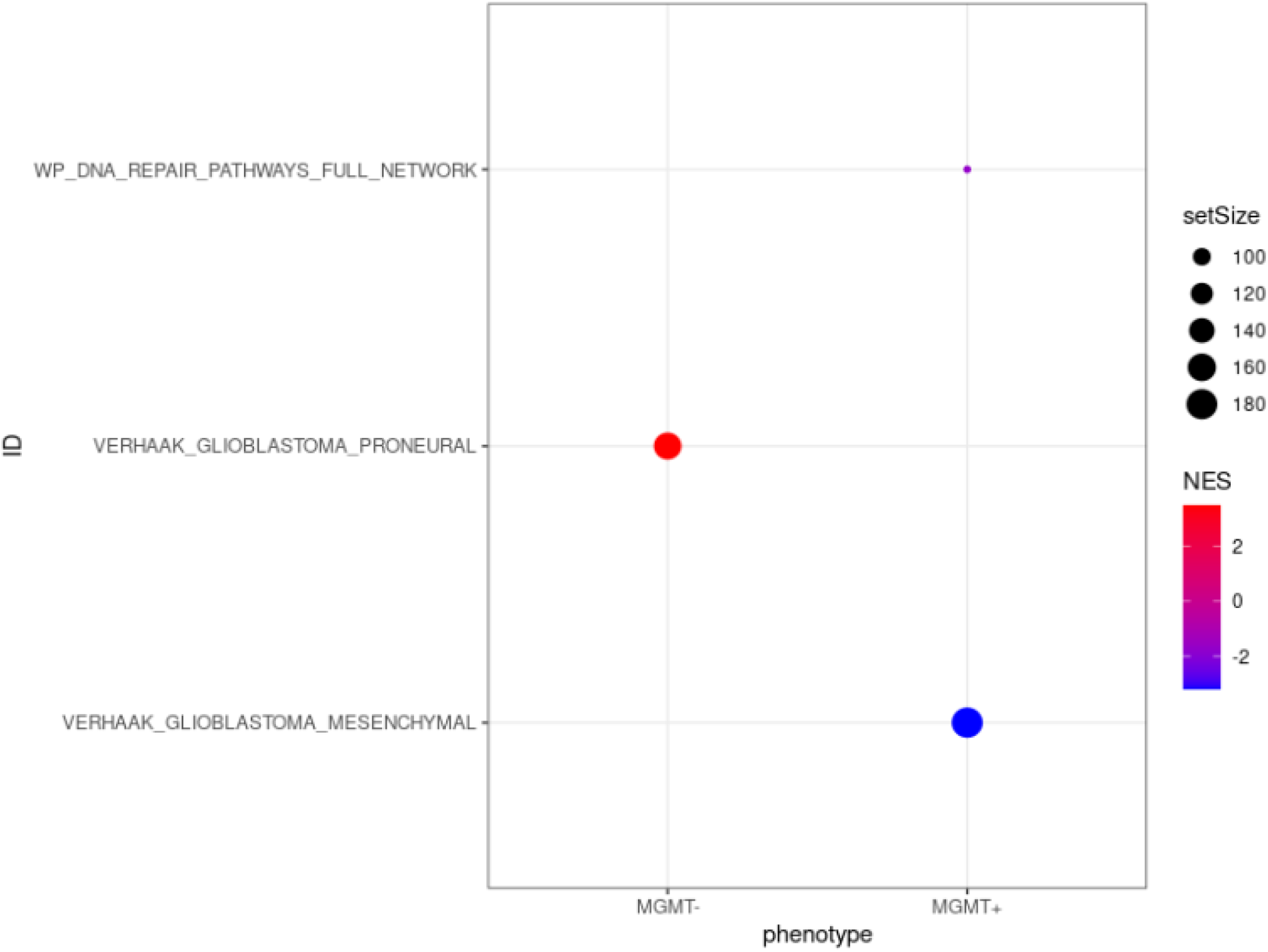
Gene set enrichment analysis showing MGMT+ cells are enriched with DNA repair pathways and Verhaak mesenchymal signature, whereas MGMT-cells are enriched with Verhaak proneural signature

### Cell cycling per MGMT expression

By applying cell-cycle scores to the CPTAC cohort per Seurat methods, 80.6% of MGMT+ cells were non-cycling whereas 56.8% of MGMT-cells were non-cycling. The difference is smaller in the Neftel et al study: 63.7% of MGMT+ cells were non-cycling and 69.98% of MGMT-cells were non-cycling.

### Find differentially expressed markers between MGMT+ and MGMT-cells

We then used Seurat’s FindMarkers function to find differentially expressed genes between cells expressing MGMT and cells not expressing MGMT. We used the unmethylated cases in the CPTAC cohort for this analysis as the unmethylated cases include a higher proportion of MGMT expressing cells. Supplementary table 1 includes the results of this analysis.

### Functional enrichment analysis of differentially expressed genes

Gene Set Enrichment Analysis (GSEA) of the differentially expressed genes between MGMT+ and MGMT-cells was applied. Interestingly, the VERHAAK_GLIOBLASTOMA_PRONEURAL set had the highest normalized enrichment score (NES) in the MGMT negative cells (NES 3.49, adjusted p value 2.173636e-08). On the other hand, the VERHAAK_GLIOBLASTOMA_ MESENCHYMAL set had the highest NES in the MGMT+ cells (NES -3.2, adjusted p value 2.173636e-08). The WP_DNA_REPAIR_PATHWAYS_FULL_NETWORK (genes: MGMT, POLE4, FANCF, CETN2, DDB2) set was enriched in the MGMT+ cells (NES -1.71, adjusted p value 0.042) (Figure 3).

## Discussion

The MGMT promoter methylation status is the most established predictive marker of response to temozolomide in GBM. In this manuscript, we aimed to explore MGMT expression at the single-cell level considering MGMTp methylation status. We used publicly available data from 3 scRNA/snRNA sequencing studies. The CPTAC cohort is well annotated and includes information about MGMT promoter methylation status for samples as determined by the MGMT-STP27 model from DNA methylation data. In the CPTAC cohort, the median expression of MGMT was 0.82% in the MGMTp methylated group and 5.7% in the unmethylated group. MGMT expression was <2% in MGMTp methylated group. Therefore 2% can be used in scRNA/snRNA experiments to predict the MGMTp methylation status of the sample.

In the Neftel study, the median expression of MGMT was 0.59% in the MGMTp methylated group and 12.44% in the MGMT. Three MGMTp unmethylated samples in the Neftel cohort did not follow the 2% rule. It is unclear if this is due to simply inaccurate annotation of these samples. Moreover, the Neftel paper did not specify the method used to detect MGMTp methylation. Or could it be that truly MGMTp unmethylated samples can have a low MGMT expression status? Could this explain why some unmethylated MGMTp patients surpass the expected survival? Bigger longitudinal studies that correlate MGMT expression based on scRNA data with survival are needed to determine the prognostic significance of MGMT expression on survival.

The possibility of MGMTp methylation status to change at GBM recurrence has been previously reported, more commonly from methylated to unmethylated status but also the other way around (7, 8). We confirm this finding by using the scRNA/snRNA data. By using the Wang, et al, dataset that included pairs of primary and recurrent GBM samples, and of the 21 pairs, MGMT expression decreased at recurrence in 11 pairs and increased in 10 pairs. By using the 2% cutoff, MGMTp methylation status changed from unmethylated to methylated in 4/21 pairs (19%) and from methylated to unmethylated in 4/21 pairs (19%).

We then identified differentially expressed genes between MGMT expressing and MGMT negative cells. Functional enrichment analysis using GSEA revealed that MGMT expressing cells are enriched with mesenchymal genes and MGMT negative cells are expressed with proneural genes. Unsupervised hierarchical clustering of bulk RNA data from the TCGA network recognized 3 distinct molecular IDHwt GBM subtypes: proneural, classical, and mesenchymal (15). The mesenchymal subtype has always been linked to aggressive behaviour. The fact that MGMT expressing cells are enriched with the mesenchymal subtype genes support this notion. Morever, MGMT expressing cells do not appear to be more cycling than MGMT negative cells. And markers of “stem cells” such as CD133 and CD15 (16) do not appear to be more expressed in MGMT+ cells.

## Supporting information

sessionInfo

## Supplementary material

### CPTAC (11) (PMID: 33577785)

The supplementary table “1-s2.0-S1535610821000507-mmc2” includes the clinical characteristic data and was downloaded from the original paper. The case IDs for the snRNA samples were downloaded from the Genomics Data Commons (GDC) Data Portal under the file “repository-cases-table.2024-04-10.tsv”. The barcodes, features and matrices files were downloaded from GDC using UUIDs from the manifest file.

snrna_merged_v2020-08-05_cell_metadata.parquet was downloaded from the NCI Proteomic Data Commons. It included data regarding the MGMT promoter methylation status and other molecular characterizations of the samples (e.g. EGFR amplification status and TERT promoter methylation status). It also included the cell-type annotation (tumor cells versus microenvironment) per the original paper’s methods.

### Neftel (9) (PMID: 31327527)

1-s2.0-S0092867419306877-mmc1 supplementary table was downloaded from the original paper and included the clinical characteristics for the cohort including the MGMT promoter methylation status (method not specified).

### sessionInfo

The output from running ‘sessionInfo’ details all the R packages and versions used in this script

### Supplementary table 1

MGMT_markers

## Code availability

All code used is available on GitHub (https://github.com/iyadalnahhas/scRNA_MGMT/blob/main/scRNA_MGMT.Rmd)

