## Supplementary material for "What scRNA sequencing taught us about MGMT expression in glioblastoma": sessionInfo

R version 4.3.1 (2023-06-16)

Platform: x86_64-pc-linux-gnu (64-bit)

Running under: Red Hat Enterprise Linux 8.10 (Ootpa)

Matrix products: default

BLAS/LAPACK: /usr/lib64/libopenblasp-r0.3.15.so; LAPACK version 3.9.0

locale:

[1] LC_CTYPE=en_US.UTF-8 LC_NUMERIC=C LC_TIME=en_US.UTF-8

[4] LC_COLLATE=en_US.UTF-8 LC_MONETARY=en_US.UTF-8 LC_MESSAGES=en_US.UTF-8

[7] LC_PAPER=en_US.UTF-8 LC_NAME=C LC_ADDRESS=C

[10] LC_TELEPHONE=C LC_MEASUREMENT=en_US.UTF-8 LC_IDENTIFICATION=C

time zone: America/New_York

tzcode source: system (glibc)

attached base packages:

[1] stats graphics grDevices utils datasets methods base

loaded via a namespace (and not attached):

[1] RColorBrewer_1.1-3 rstudioapi_0.16.0 jsonlite_1.8.8

[4] magrittr_2.0.3 spatstat.utils_3.0-3 rmarkdown_2.23

[7] fs_1.6.2 zlibbioc_1.48.2 vctrs_0.6.5

[10] ROCR_1.0-11 memoise_2.0.1 spatstat.explore_3.2-1

[13] RCurl_1.98-1.12 htmltools_0.5.5 sctransform_0.4.1

```
[16] parallelly_1.36.0      KernSmooth_2.23-21     htmlwidgets_1.6.2
[19] ica_1.0-3              plyr_1.8.8             plotly_4.10.4
[22] zoo_1.8-12             cachem_1.0.8           igraph_1.5.0
[25] mime_0.12              lifecycle_1.0.4        pkgconfig_2.0.3
[28] Matrix_1.6-5           R6_2.5.1               fastmap_1.1.1
[31] GenomeInfoDbData_1.2.11 fitdistrplus_1.1-11    future_1.33.0
[34] shiny_1.7.4.1          digest_0.6.33          colorspace_2.1-0
[37] patchwork_1.2.0.9000   AnnotationDbi_1.64.1   S4Vectors_0.40.2
[40] tensor_1.5             Seurat_5.0.3           RSpectra_0.16-1
[43] irlba_2.3.5.1          RSQLite_2.3.1          progressr_0.13.0
[46] spatstat.sparse_3.0-2  fansi_1.0.6            polyclip_1.10-4
[49] abind_1.4-5            httr_1.4.6             compiler_4.3.1
[52] bit64_4.0.5            BiocParallel_1.36.0    DBI_1.1.3
[55] fastDummies_1.7.3      MASS_7.3-60            HDO.db_0.99.1
[58] tools_4.3.1            lmtest_0.9-40          httpuv_1.6.11
[61] future.apply_1.11.0    goftest_1.2-3          glue_1.7.0
[64] nlme_3.1-162           GOSemSim_2.28.1        promises_1.2.0.1
[67] grid_4.3.1             Rtsne_0.17             cluster_2.1.4
[70] reshape2_1.4.4         fgsea_1.28.0           generics_0.1.3
[73] spatstat.data_3.0-1    gtable_0.3.5           tidyr_1.3.0
[76] data.table_1.14.8      sp_2.0-0               utf8_1.2.4
[79] XVector_0.42.0         spatstat.geom_3.2-2    BiocGenerics_0.48.1
[82] RcppAnnoy_0.0.21       ggrepel_0.9.3          RANN_2.6.1
[85] pillar_1.9.0           stringr_1.5.0          yulab.utils_0.1.4
[88] spam_2.9-1             RcppHNSW_0.4.1         later_1.3.1
[91] splines_4.3.1          dplyr_1.1.2            lattice_0.21-8
```

```
 [94] deldir_1.0-9          survival_3.5-5         bit_4.0.5
 [97] tidyselect_1.2.1      GO.db_3.18.0           Biostrings_2.70.3
[100] miniUI_0.1.1.1        pbapply_1.7-2          knitr_1.43
[103] gridExtra_2.3         IRanges_2.36.0         scattermore_1.2
[106] xfun_0.39             stats4_4.3.1           Biobase_2.62.0
[109] matrixStats_1.0.0     stringi_1.7.12         yaml_2.3.8
[112] lazyeval_0.2.2        evaluate_0.21          codetools_0.2-19
[115] tibble_3.2.1          qvalue_2.34.0          cli_3.6.2
[118] uwot_0.1.16           xtable_1.8-4           reticulate_1.36.1
[121] munsell_0.5.1         Rcpp_1.0.12            GenomeInfoDb_1.38.8
[124] spatstat.random_3.1-5 globals_0.16.2         png_0.1-8
[127] parallel_4.3.1        ellipsis_0.3.2         ggplot2_3.5.0.9000
[130] blob_1.2.4            dotCall64_1.0-2        DOSE_3.28.2
[133] bitops_1.0-7          listenv_0.9.0          viridisLite_0.4.2
[136] scales_1.3.0          ggridges_0.5.4         SeuratObject_5.0.1
[139] leiden_0.4.3          purrr_1.0.1            crayon_1.5.2
[142] rlang_1.1.3           cowplot_1.1.1          fastmatch_1.1-4
[145] KEGGREST_1.42.0
```
